# Myeloid-derived alveolar-like macrophages are a tractable model to understand the role of ontogeny in alveolar macrophage function *ex vivo* and in the lungs

**DOI:** 10.64898/2026.05.19.726293

**Authors:** Reham A. Ammar, Andrew J. Olive

## Abstract

Alveolar macrophages (AMs) are tissue-resident and the primary immune cells in the airspace. Following perturbations in the lungs, these AMs that are derived from the fetal liver, become depleted and are transiently replaced by myeloid cells that use lung-specific cues to differentiate into myeloid-derived AMs. While these myeloid-derived AMs are critically important in a range of pulmonary diseases, including post-influenza bacterial pneumonia, it remains challenging to fully understand their function due to a lack of *ex vivo* models that recapitulate key differences observed *in vivo* between AMs and myeloid-derived AMs. Here, we overcome this limitation by expanding our recently developed model of fetal liver-derived alveolar macrophages (FLAMs) to differentiate myeloid progenitors in the presence of GM-CSF and TGFβ, key cytokines that drive tissue resident AM functions. These myeloid-derived alveolar-like macrophages (MAMs) express AM surface markers and look similar morphologically to FLAMs, however, they remain more inflammatory than FLAMs. Mechanistic studies found that differential CpG methylation at inflammatory loci, basal transcriptional expression, and metabolic flux all contribute to the hyperinflammatory state of MAMs. Importantly, we find that while FLAMs are highly dependent of lipid metabolism, MAMs are more glycolytic and this hardwired metabolism is not easily overcome to mute their inflammatory state. Finally, we found that MAMs and FLAMs both function within the lung environment following transfer into mice lacking AMs. While both MAMs and FLAMs stably seed the lungs and reverse pulmonary proteinosis, MAMs remain highly inflammatory in the lungs following an LPS model of acute lung injury. Taken together our results find that MAMs are a reproducible model of myeloid-derived AMs and lays the groundwork to better understand how these important immune cells contribute to pulmonary homeostasis and responses to lung perturbations. These future studies will help to identify new targets that can be modulated to prevent severe pulmonary disease outcomes.

## Introduction

Positioned at the interface between the external environment and internal milieu, the lungs are a key battleground for pathogens and environmental toxicants. Consistent exposure to these external stimuli necessitates a fine-tuned immune response to curb potential respiratory disease while maintaining lung physiological functions. Lung immune cells like macrophages play a key role in regulating lung immune responses^1,2^. Lung macrophage populations can originate from either the fetal liver or myeloid progenitor cells. The dynamic effector functions of these macrophages are dependent on their developmental origin. Tissue resident alveolar macrophages (**AMs**) develop from fetal livers during early embryonic stages. They populate the alveolar space and self-renew during their lifetime. AMs maintain lung homeostasis by balancing inflammation during infections or exposure to particulate matter^3,4^. The lung microenvironment produces cytokines including Granulocyte-macrophage colony-stimulating factor (**GM-CSF**) and Transforming growth factor-beta (**TGFβ**) that regulate the inflammatory response of AMs ^5–9^. However, during acute lung infections or inflammation, AMs are depleted^10^. To compensate for the loss of AMs, myeloid-derived monocytes are recruited to the alveoli. Local signals in the alveolar space instruct the differentiation of these recruited monocytes into mature myeloid-derived alveolar macrophages, yet data suggest that these recruited myeloid-derived alveolar macrophages remain functionally distinct, especially regarding inflammatory regulation^11–13^. Dysregulation of the lung macrophages contributes to important lung disease conditions including acute lung injury and post-influenza bacterial pneumonia^14^. Unfortunately, we currently lack a full understanding of the mechanisms underlying this heterogeneous immune response due to limited models that recapitulate distinct macrophage subsets in the lungs. A greater understanding of distinct lung macrophage inflammatory responses is imperative to uncover pathways that may contribute to susceptibility to infections and inflammatory disease in the pulmonary tissue and ultimately help identify novel therapeutic targets for these conditions.

One key pathway in pulmonary inflammation is the activation of the pattern recognition receptor toll-like receptor-4 (TLR4)^15^. While TLR4, is activated by a wide range of pathogen associated molecular patterns (**PAMPs**) it is an important sensor for gram-negative bacterial derived lipopolysaccharide (**LPS**)^16^. To sense LPS efficiently, LPS binding proteins coordinate with the co-receptor CD14 which increases the sensitivity of TLR4 to LPS stimulation^17^. Upon TLR4 activation, parallel signaling pathways mediated by MyD88 and TRIF activate key transcription factors in the inflammatory response including NF-kβ, MAP kinases, and interferon regulatory factors 3 and 7 (IRF3 and IRF7). These transcription factors then coordinate the induction of cytokines including TNF, IL-6, IL1, and type I IFN^18,19^. Dysregulation of these cytokines in the lungs can drive pulmonary inflammatory disease. Thus, understanding how these distinct signaling pathways downstream of TLR4 regulate pulmonary macrophage inflammatory responses to pathologic damage is critically important.

To drive robust cytokine production, LPS activation is known to modulate the cellular metabolic profile. In seminal work it was found that TLR4 binding LPS in bone marrow derived macrophages (BMDMs) results in a Warburg effect where macrophages shift away from oxidative phosphorylation to become more glycolytic ^20^. This is partially mediated through the accumulation of succinate which blocks the activity of prolyl hydroxylase, leading to the stabilization of hypoxia inducible factor 1a (HIF1α). HIF1α then contributes further to proinflammatory cytokine production^11,21^. Subsequent work showed that glycolysis is preferentially used by activated BMDMs to support the rapid ATP demand for cytokine production and the regulation of inflammatory pathways by glycolytic intermediates ^22,23^. While these studies show BMDMs respond to LPS with large metabolic shifts, BMDMs are not a representative model of tissue resident AMs or recruited myeloid-derived alveolar macrophages. BMDMs are differentiated in the cytokine M-CSF and are generally thought to model recruited inflammatory macrophages in the lungs that are distinct from other lung macrophage subsets^24^. It is important to determine the macrophage subset specific responses to be better positioned to modulate pulmonary disease therapeutically.

Recent work has shown that AMs have fundamentally distinct responses to TLR4 activation from BMDMs. AMs are less sensitive to LPS activation resulting in reduced inflammatory responses. This is partially mediated by the recalcitrance of AMs to shift their metabolism away from OXPHOS towards glycolysis. While the underlying mechanisms of the AM hypoinflammatory state remain to fully understood, one hypothesis is that these cells are highly dependent on lipid metabolism and OxPhos to ensure the homeostatic function of AMs to metabolize secreted surfactants in the lungs. AMs express high levels of the nuclear transcription factor peroxisome proliferator-activated receptor gamma (**PPARγ**) to regulate lipid metabolism, and mice deficient in PPARγ lack the presence of functional AMs ^25,26^. In addition, mitochondria and peroxisomes are closely associated and play a crucial role in regulating cellular respiration. Peroxisomes are the lipid metabolism hub in cells and play a secondary role in managing oxidative stress through the expression of catalase. Recent work by us and others showed that AMs have significantly higher numbers of peroxisomes, and these peroxisomes play an important role in maintain AM functions^27,28^. In addition, AMs express high levels of ATP Synthase highlighting the central importance of mitochondrial OXPHOS in these cells. Disruption of the AM metabolic state can result in the production of mitochondrial ROS that disrupts mitochondrial membrane potential leading to the release of mitochondrial nucleotides into the cytosol where they can activate type I IFN responses^28,29^. A key link between the mitochondria and peroxisomes is PPARγ as it regulates transcription of genes responsible for peroxisomal function and β-oxidation^25,26^. Thus, there are major differences in how BMDMs and AMs regulate their metabolism and the subsequent response to LPS activation.

Much of our understanding of inflammatory regulation comes from work using BMDMs due to a lack of models that recapitulate key functions of AMs and myeloid-derived AMs. Recent work by us and others have overcome this limitation by developing new *ex vivo* models of AMs ^25,30^. In our model which we call fetal liver derived alveolar like macrophages (**FLAM**s), fetal liver cells are grown in the presence of the lung cytokines GM-CSF and TGFΒ resulting in a long-term stable population that recapitulate key AM functions and phenotypes ^28,31^. This model is useful to characterize key signaling networks in AMs as it is genetically tractable and cells can be propagated, frozen, and revived without any immortalization. While FLAMs enable us to model the tissue resident AM population, we still lacked tractable models to understand differences in myeloid-derived alveolar macrophages, leaving major gaps in our understanding of this important lung macrophage subset.

Here, we introduce a tractable model of myeloid-derived AMs. We find that myeloid progenitors differentiated in GM-CSF and TGFΒ, which we call myeloid-derived alveolar-like macrophages (MAMs), results in a population of cells that are transcriptionally similar to AMs at baseline and express key AM markers including SiglecF and CD11a. However, we find that MAMs are much more inflammatory to the same concentrations LPS compared to FLAMs, recapitulating critical phenotypes observed in lungs. Mechanistic studies found distinct epigenetic methylation profiles between MAMs in FLAMs particularly near inflammatory loci. Metabolic studies also found that MAMs are highly glycolytic compared to FLAMs and this hardwired metabolic state cannot be easily disrupted by modulating lipid metabolism. Thus, we can use MAMs as tool to understand the key regulatory differences between myeloid-derived AMs and tissue resident AMs. Finally, to test the appropriateness of our model, we optimized *in vivo* transfer studies to examine how MAMs and FLAMs function within the lung environment. In line with our *ex vivo* results we found that lungs seeded with MAMs result in more severe inflammatory responses following intranasal LPS activation compared to FLAMs. Collectively, our data show that MAMs are tractable model of myeloid-derived AMs that can be studied both *ex vivo* and *in vivo* and highlights the key role ontogeny plays in dictating pulmonary inflammation.

## Materials and Methods

### Animals

Experimental protocols were approved by the Institutional Animal Care and Use Committees at Michigan State University (animal use form [AUF] no. PROTO202500031). All protocols were strictly adhered to throughout the entire study. Six- to 8-wk-old C57BL/6J mice (catalog no. 000664), were obtained from The Jackson Laboratory (Bar Harbor, ME). Mice were given free access to food and water under controlled conditions (humidity, 40–55%; lighting, 12-hour light/12-hour dark cycles; and temperature, 24 ± 2°C), as described previously. (32). Pregnant dams at 8–10 weeks of age and 14–18 gestational days were euthanized to obtain murine fetuses to generate FLAMs. MAMs and BMDMs were isolated from male and female mice >10 weeks of age.

### FLAMs, MAMs, BMDMs isolation and culture

FLAMs were developed from fetal livers that were isolated from pregnant C57B6L/J mice after euthanasia with CO2 followed by cervical dislocation. FLAMs were then cultured in RPMI media with 10% fetal bovine serum (FBS), 20 ng/ml TGFβ and 30 ng/ml GM-CSF as described before. Primary MAMs were developed from myeloid progenitors isolated from C57B6L/J bone marrow and then differentiated into mature AM-like cells in RMPI media containing TGFβ and GM-CSF similar to FLAMs. Primary BMDMs were isolated similar to MAMs but differentiated into mature macrophages in DMEM media containing 10%FBS and 25 ng/ml recombinant M-CSF. Cell cultures are maintained in 5 µg/ml Ciprofloxacin for the first 7 days. Cells were incubated at 37C in 5% CO2 and were regularly assessed.

### TLRs activation

For macrophages immune activation, cells were treated with 12.5 ng/ml ultrapure LPS (TLR4 agonist, Thermofisher, Cat#tlrl-pb5lps) at indicated timepoints.

### Cytokine analysis

At the indicated time points the culture supernatants or bronchial lavage fluid were collected and assessed for the expression of cytokine levels including IFNβ using LumiKine™ Xpress mIFN-β Kit (Cat no: luex-mIFNβv3), IL6, TNFa, IL1a, CXCL10 (using DuoSet R&D systems ELISA kits) according to the manufacturer instructions. Using a Spark multimode plate reader (Tecan), absorbance at λ=450nm was measured.

### Flow cytometry

Plated cells were lifted at the indicated time points and stained with Zombie live/dead stain (Biolegend) for 15 mins. Cells then were washed twice with PBS and stained with fluorescent antibody mixture as indicated for 20 mins including anti-CD170 (SiglecF), Siglec1, CD11c, CD11b, CD11a, CD14, MHCII, TLR4, TLR2 at 1:400 dilution in PBS (Biolegend). Afterwards, cells were washed three times with PBS and fixed with 1% paraformaldehyde (Electron Microscopy Sciences) for 15 mins. Flow cytometry was performed on an Attune Cytpix analyzer (Thermofischer scientific) at the Michigan State University flowcytometry core. Data analysis was done using FlowJo software 10.10.0.

### Rosiglitazone, Infasurf, Oxamate treatments

Cells were treated with 20 µM Rosiglitazone (Sigma Aldrich) overnight, or 20mM Oxamate (Thermofischer Scientific) for 1 hour then stimulated with 12.5 ng/ml LPS for the indicated time points (6, 16, 24 hours). For Infasurf (ONY Biotech), cells were maintained in a culture media containing 100 µg/ml infasurf then stimulated with 12.5 ng/ml LPS for the indicated time points.

### RNA extraction

Cells are TRIzol reagent (life technologies) was added to each well and incubated at room temperature for 5 minutes to allow for cell lysis. Cell lysate was transferred to 1.5 ml tubes and 100 µl of chloroform reagent was added, vigorously mixed, and centrifuged at 10,000xg for 18 minutes at 4C to separate mRNA. Afterwards, the aqueous layer was separated and mixed with equal volume of absolute ethanol. The mixture was then processed with Qiagen RNA extraction kit according to manufacturer instructions. RNA was assessed for quantity and purity using a NanoDrop. Amplification of RNA was done according to manufacturer instructions using the Qiagen one-step Syber Green RT-PCR kit. Relative expression levels of mRNA were calculated after normalization to GAPDH/β-actin.

### RNA-sequencing analysis

We used the Direct-zol RNA Extraction Kit (Zymo Research, Cat no. R2072) to extract RNA according to the manufacturer’s protocol. The Illumina Stranded mRNA Library Prep kit (Illumina, Cat no. 20040534) with IDT for Illumina RNA Unique Dual Index adapters was used for library preparation following the manufacturer’s recommendations but using half-volume reactions. Qubit™ dsDNA HS (ThermoFischer Scientific, Cat no. Q32851) and Agilent 4200 TapeStation HS DNA1000 assays (Agilent, Cat no. 5067-5584) were used to measure quality and quantity of the generated libraries. The libraries were pooled in equimolar amounts, and the Invitrogen Collibri Quantification qPCR kit (Invitrogen, Cat no. A38524100) was used to quantify the pooled library. The pool was loaded onto 2 lanes of a NovaSeq S4 flow cell, and sequencing was performed in a 2×150 bp paired end format using a NovaSeq 6000 v1.5 100-cycle reagent kit (Illumina, Cat no. 20028316). Base calling was performed with Illumina Real Time Analysis (RTA; Version 3.4.4), and the output of RTA was demultiplexed and converted to the FastQ format with Illumina Bcl2fastq (Version 2.20.0).

A Snakemake pipeline version 7.32.4 (https://github.com/kaylaconner/olivelab-rnaseq/tree/main) was built to run RNA reads quality assessment, mapping, and counting through the Michigan State University’s Institute for Cyber-Enabled Research. FastQC version 0.12.1 was used to assess reads quality. Bowtie 2 version 2.5.1 was used for RNA reads alignment against the GRCm39 mouse reference genome then the featureCounts function (subread package version 2.0.6) was used to assess the aligned reads. Differential gene expression analysis in MAMs compared to FLAMs under resting and activated conditions was run through DESeq2 package (version 1.50.1) in R (version 4.5.2). A cut off value of 10 reads in three samples was used to filter out genes with low read number. Pathway analysis was performed through GSEA (version 4.3.3) on a pre-ranked DESeq2 gene list for mouse hallmark pathways (https://www.gseamsigdb.org/gsea/index.jsp)

For data visualization, Heatmaps were generated for the DESeq2 normalized counts in R (version 4.5.2) using pheatmap package (version1.0.13), Principal Component Analysis (PCA) was run using R on genes normalized counts matrices. ggplot2 (version 4.0.1) was used for PCA visualization through the autoplot function.

### Infinium mouse methylation BeadChip assay

Genomic DNA was extracted for FLAMs and MAMs under resting and LPS-treated conditions using the DNAeasy kit (Qiagen, USA, Cat# 69506). DNA was quantified and checked for quality on a Qubit fluorometer using a Qubit dsDNA quantification assay kit (Thermofisher scientific, Cat# Q32853). 500ng of high-quality gDNA in 45ul 10mM Tris-buffer (Fisherscientific, Cat# AM9855G) was stored in -80C and submitted to Van Andel Institute genomics core, (Grand Rapids, Michigan) in three biological replicates to run the Illumina infinium mouse methylation BeadChip assay targeting 285K CpG sites including genes, promoters, and CpG islands. Data import was run using RnBeads package (version 2.28.0) in R (version 2025.09.2) for mouse probe and read intensity. A preprocessing step was done to filter out any probes without intensity value, sex chromosome probes, off-target probes. Probe annotation was done with the RnBeads mm10 package (version 2.18.0). Data normalization was done using the bmiq function while background correction was done using the methylumi.noob function withing the RnBeads package. The ratio between methylated probe intensity to the total intensity (methylated + unmethylated probe intensities) was calculated as β-value. A β-value of 0 indicated no methylation while a β-value of 1 indicated complete methylation. Differential methylation analysis was done using the ComputeDiffMeth function within the RnBeads package to calculate the Δβ values at the indicated pairwise group comparisons. A cutoff value of 0.05 was set for the adjusted *P value* for all differential methylation data. Gene ontology (GO) analysis was performed using Metascape ( <u>Zhou et al., Nature Communication (2019), 10(1):1523</u>; [http://metascape.org]) for the differentially hypomethylated and hypermethylated CpG sites associated with genes and promotor regions.

### Extracellular metabolic flux assays

Real-time oxygen consumption rate (OCR) and extracellular acidification rate (ECAR) was measured using a Seahorse XFe96 Analyzer according to the manufacturer instructions (Agilent Technologies). Seahorse media was prepared using XF RPMI medium (Cat#103576-100), Glucose Solution (cat#103577-100), Pyruvate Solution (Cat#103578-100), Glutamine Solution (Cat#103579-100). To measure mitochondrial function, Mito stress test was performed through the sequential addition of 10 μM oligomycin (Thermofisher, Cat# J61898), 15 μM FCCP (Sigma, Cat# C2920), and 10uM rotenone (Sigma, R8875-1G) and antimycin A (Sigma, Cat#A8674). To assess the glycolytic function, glycolysis stress test was performed to measure ECAR after the sequential addition of 10mM glucose, 10 μM oligomycin, 50 mM 2-Deoxy glucose (2-DG) (Cat#L07338.14).

### Adoptive transfer of FLAMs and MAMs into GMCSFR-KO mice

Mice lacking GMCSFR-KO were selected for experiments following weaning at 3-4 weeks prior to the development of severe pulmonary alveolar proteinosis (PAPs). We transferred 10^6^ cells/mouse in 100 ul PBS on two consecutive days intratracheally using the tongue pull method as described before ^32^ Mice were left to two weeks to allow the cells enough time to settle in the lung environment. Afterwards, 1mg/kg LPS was instilled intratracheally and incubated for 24 hours. BALF was extracted from the mice, spun down at 500xg for 5 mins and supernatants were used to measure optical density (OD600) to check for the turbidity as well as to measure inflammatory cytokines with ELISAs. Flow cytometry was used to measure mean fluorescence intensity of AM/neutrophile-specific surface markers expression on the cellular component of the BALF.

### Scanning electron microscopy

2.5 x 10^5^ MAMs or FLAMs were seeded on glass circular coverslips (Electron Microscopy Sciences) in 100uL cytokine media. Each coverslip was placed in a separate well of a 6-well plate. Cells were allowed to attach for 5 mins then 1 ml of media was added on top. Cells were then fixed by 4% glutaraldehyde buffered with 0.01M sodium phosphate for 30 minutes. A gradual dehydration step was performed using serial concentrations of ethanol (25, 50, 75, 95%) for 10 minutes incubation with each concentration followed by 10 minutes changes in 100% ethanol.

Each sample was critical point dried in a Leica Microsystems model EM CPD300 critical point dryer (Leica Microsystems, Vienna, Austria) using carbon dioxide as the transitional fluid. Afterwards, samples were mounted on aluminum subs using epoxy glue (System Three Quick Cure 5, System Three Resins, Inc., Auburn, WA). Finally, samples were coated with osmium at 10 nm thickness in a Tennant20 osmium CVD (chemical vapor deposition) coater (Meiwafosis Co., Ltd., Osaka, Japan). For imaging, a JEOL 7500F (field emission emitter) scanning electron microscope (JEOL Ltd., Tokyo, Japan) was used.

### Statistical analysis and data visualization

Statistical analysis was performed using GraphPad prism version 10.6.0. Parametric data sets were analyzed using either student-t test to compare 2 groups, or one-way/two-way analysis of variance (ANOVA) tests to compare >2 groups followed by Tukey-Kramer post hoc multiple comparison statistical tests. Unless otherwise stated, data are presented as mean ± standard deviation.

### Data accessibility

All processed data is included in the manuscript. Raw unprocessed data is being deposited into appropriate databases and accession numbers will be added.

## Results

### MAMs express AM-specific markers similarly to FLAMs while exhibiting a differential inflammatory profile

We previously developed FLAMs as an *ex vivo* model for tissue-resident alveolar macrophages. For this model fetal liver cells are grown in the presence of GM-CSF and TGFΒ. Since we lack useful models of myeloid-derived alveolar macrophages, we tested the hypothesis that differentiating myeloid-progenitors in media containing GM-CSF and TGFΒ would generate myeloid-derived AM-like cells (MAMs) (Figure 1A). To test this hypothesis, we first compared the cellular morphology of FLAMs and MAMs by scanning electron microscopy (SEM). We observed that both MAMs and FLAMs had similar circular morphology with ruffled surface structures (Figure 1B). We next examined the surface markers associated with AMs, inflammatory macrophages, and TLR4 (Figure 1C). BMDMs expressed high levels of CD14 and CD11b while expressing almost no SiglecF and CD11a. In contrast, MAMs looked very similar to FLAMs with high levels of SiglecF and CD11a and low levels of CD14 and CD11b. Importantly, MAMs, and FLAMs expressed similar surface levels of TLR4. These data suggest that differentiating myeloid progenitors in FLAM media drives a phenotypic state that resembles AMs.

**Figure 1.**
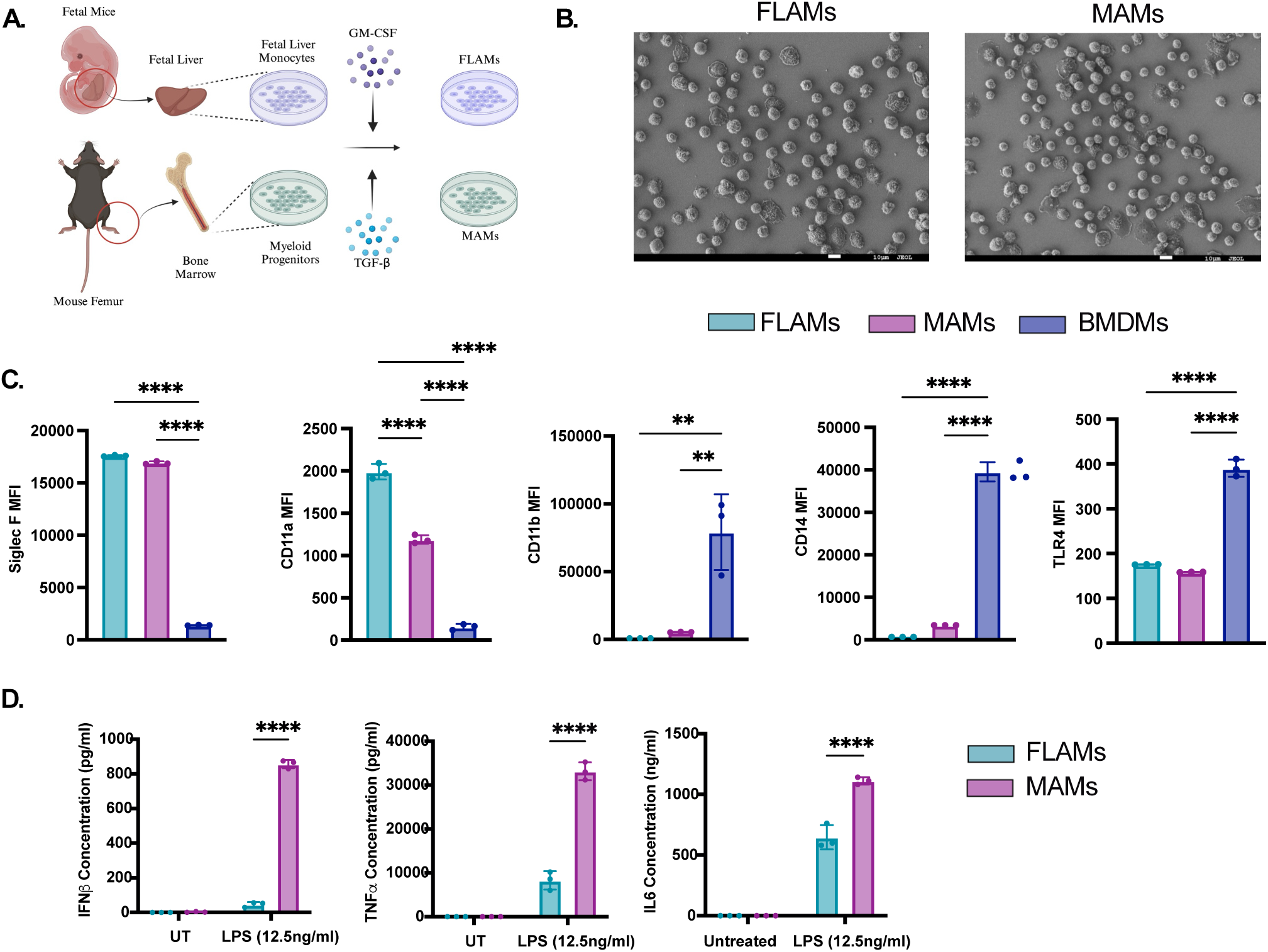
MAMs resemble myeloid-derived AMs and are more inflammatory than FLAMs. **A)** Schematic representation of methods to differentiate FLAMs and MAMs. Figure made using Biorender. **B)** FLAMs and MAMs were imaged using scanning electron microscopy. Shown is a representative image from two independent samples. Bars indicate 10uM. **C)** The expression of the indicated surface markers was quantified by flow cytometry from FLAMs, MAMs, and BMDMs. Shown is the quantified mean fluorescence intensity. Each point represents a technical replicate from one representative experiment of 3. **p<0.01 by one-way ANOVA with tukey test. **D)** FLAMs, MAMs, and BMDMs were stimulated with LPS (12.5ng/ml) for 24 hrs. Supernatants were collected for quantification of IFNβ, TNF, and IL6 by ELISA. Shown is one representative experiment of three each containing 3 replicates per experiment. **p<0.01 ***p<.001 by one-way ANOVA with a Tukey post-hoc test for multiple comparisons.

In the lungs, a key difference between tissue resident AMs and myeloid-derived AMs is the inflammatory response, with myeloid-derived AMs driving more robust cytokine induction. To test whether MAMs and FLAMs recapitulate these key *in vivo* observations, we next stimulated FLAMs and MAMs with LPS and measured the production of the inflammatory cytokines TNF, IL6 and IFNβ (Figure 1D). In line with published *in vivo* observations, we found that MAMs induced significantly more of each cytokine compared to FLAMs. Since both MAMs and FLAMs are genetically identical and they are cultured in identical media containing GMCSF and TGFΒ, these data suggest that ontologically driven differences play a key role in regulating inflammatory responses in FLAMs and MAMs.

### Differential DNA methylation at inflammatory loci is associated with the inflammatory potential of MAMs and FLAMs

We next wanted to better understand the underlying differences between MAMs and FLAMs that drive the divergent response to LPS. Previous studies suggest that ontogeny may drive differences in the epigenetic landscape of the immune cells which in turn regulates their inflammatory profiles. To test this directly in our model we quantified the methylation of CpG sites across the genome in MAMs and FLAMs that were left untreated or were treated with LPS for 2 hours (Supplementary Table 1). Principal component analysis (**PCA**) showed strong separation between FLAMs and MAMs at CpG sites in protein coding regions yet showed few changes following LPS activation (Figure 2A). We found 911 differentially methylated protein coding CpG sites between FLAMs and MAMs under resting condition among which 261 sites were hypomethylated and 650 sites were hypermethylated (adjusted *P<0.05*). Under LPS-stimulated condition, we found 970 differentially methylated protein coding CpG site between FLAMs and MAMs, of these 300 sites were hypomethylated and 679 sites were hypermethylated (adjusted *P<0.05*). We found no significant differential methylation patterns between untreated and LPS-treated condition either in FLAMs or MAMs groups (Figure 2B). Gene ontology (GO) analysis of the differentially methylated genes showed enrichment of inflammatory responses pathways including innate immunity, inflammatory cytokines, and immunometabolism (Figure 2C). We observed that type-I interferon signature was one of the most pathways including irf4, Ifit3, Ifitim3, and Rsad2 in addition to Il6 signaling being hypermethylated in FLAMs compared to MAMs with a higher Δβ value (Figure 2D). In contrast we observed hypomethylation in FLAMs for key AM genes including MARCO, a known AM surface marker. These data suggest one mechanism by which MAMs drive more robust inflammatory responses is the accessibility of promoters associated with inflammatory gene expression.

**Figure 2.**
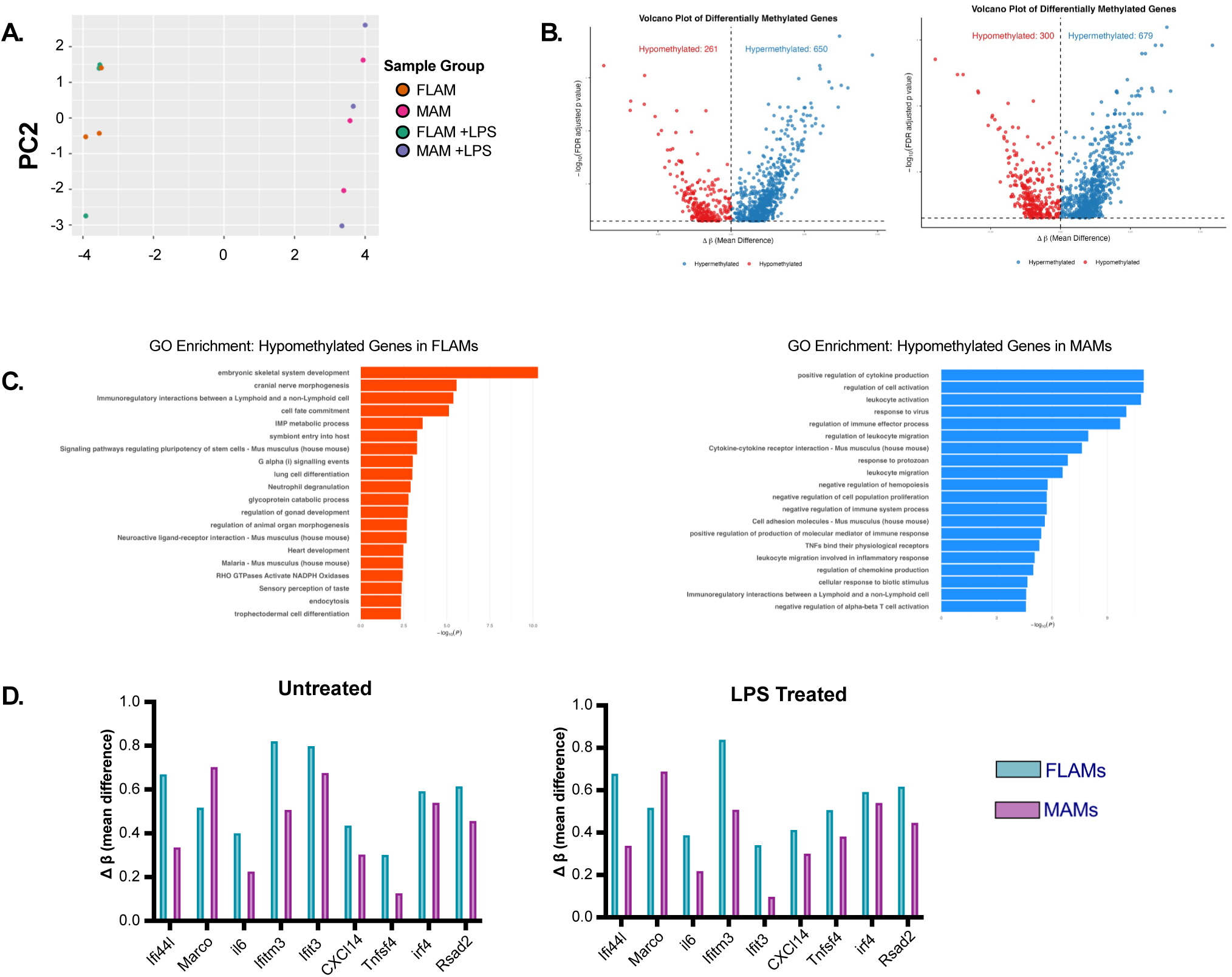
MAMs are hypomethylated at inflammatory loci compared to FLAMs. **A)** A principal component analysis (PCA) plot comparing the CpG methylation of protein expressing genes from MAMs and FLAMs that were untreated or treated with LPS (12.5ng/ml) for 24 hours. **B)** Shown is a volcano plot showing genes that are hypomethylated (Red dots) or hypermethylated (Blue dots) in FLAMs compared to MAMs that were (Left) untreated or (Right) treated with LPS (12.5ng/ml) for 24 hours. **C)** GO enrichment of pathways enriched in either hypomethylated protein coding genes in (Left) FLAMs or (Right) MAMs was determined using metascape. Data are representative of 3 biological replicates from one experiment. **D)** Shown are β-values for genes associated with IFN response and MARCO a known AM marker. Data are from three independent replicates.

### Increased inflammatory transcriptional responses are observed in MAMs following LPS treatment

While our data suggest key differences in the epigenetic landscape between MAMs and FLAMs, we next wanted to gain a more global view of transcriptional differences. To address this question, we conducted bulk RNA sequencing comparing resting MAMs and FLAMs with cells activated with LPS for six hours (Supplementary Table 2). Principal component analysis (PCA) showed resting MAMs and FLAMs cluster near each other suggesting they have similar transcriptional profiles at baseline (Figure 3A). In contrast following LPS activation, MAMs and FLAMs diverge suggesting they uniquely respond to the identical activation conditions. Among the top genes that were highly expressed in in MAMs compared to FLAMs at baseline were genes associated with antigen presentation in the major histocompatibility complex II pathway. Differential expression analysis also found genes associated with inflammation including Cxcl3 were upregulated in resting MAMs compared to FLAMs at baseline (Figure 3B). These data suggest underlying differences in how MAMs and FLAMs basally regulate their gene expression. When comparing MAMs and FLAMs activated with LPS we noted significantly higher expression of other inflammatory genes in MAMs (Figure 3B) suggesting more dramatic differences between these cells following activation. Gene set enrichment analysis (GSEA) further supports this conclusion. We observed a strong enrichment in MAMs for hallmark IFN responses and inflammatory responses (Figure 3D) in line with our initial observations. Examination of a subset of genes associated with these pathways found robust expression of cytokines, co-stimulatory molecules, and antigen presentation machinery in MAMs compared to FLAMs following LPS stimulation (Supplementary Table 2).

**Figure 3.**
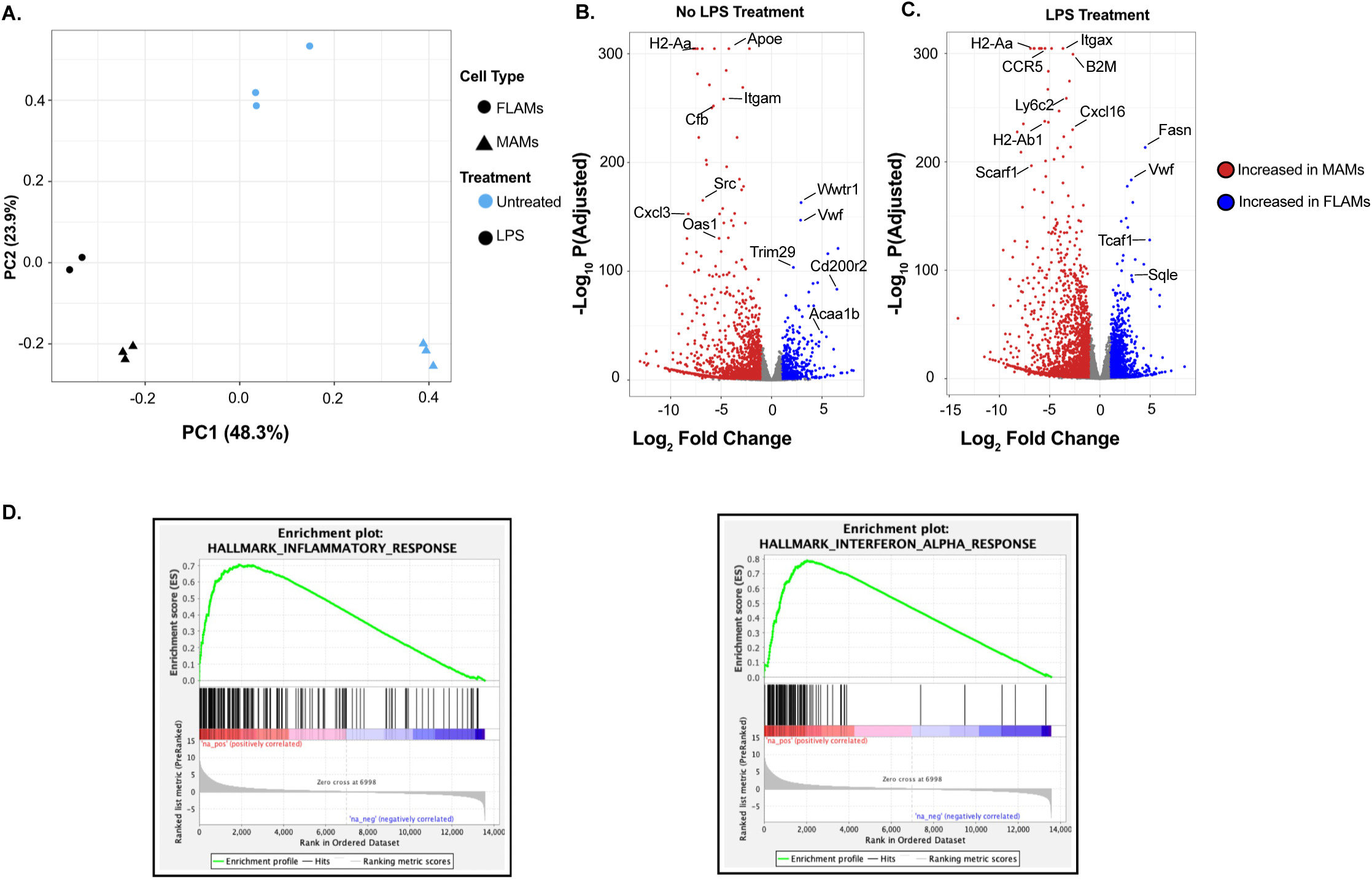
MAMs increased inflammatory response is reflected in broad transcriptional changes following LPS treatment. **A)** A principal component analysis (PCA) plot comparing the transcriptome of from MAMs and FLAMs that were untreated or treated with LPS (12.5ng/ml) for 6 hours. **B)** Shown is a volcano plot showing genes that are differentially expressed more highly in MAMs (Red dots) and FLAMs (Blue dots) in cells that were (Left) untreated or (Right) treated with LPS (12.5ng/ml) for 6 hours. **C)** Leading edge analysis of differentially expressed genes identified the IFN-alpha response and the inflammatory response hallmark pathways comparing MAMs and FLAMs treated with LPS (12.5ng/ml) for 6 hours.

### MAMs are highly glycolytic which contributes to their inflammatory state

It is well appreciated that changes in metabolism can directly impact the inflammatory state of macrophages. To directly examine how the expression of metabolic networks is regulated in MAMs in FLAMs we compared the expression of genes associated with the KEGG glycolysis and OxPhos pathways (Figure 4A). We found that following LPS stimulation, MAMs robustly induce glycolysis genes while FLAMs do not. In contrast, following LPS drove a major decrease in OxPhos genes in MAMs while FLAMs maintained high expression. When we examined a subset of genes associated with shifts between OxPhos and glycolysis we saw FLAMs maintained high expression of Idh2 and Idh3b following LPS treatment while MAMs did not (Figure 4B). Additionally, MAMs robustly induced HIF1a and Nos2 far beyond FLAMs. For Nos2 we observed over 1000-fold difference in expression between MAMs and FLAMs after LPS treatment. These data suggest metabolic differences between MAMs and FLAMs following LPS activation with FLAMs maintaining a strong OXPHOS profile while MAMs are highly glycolytic.

**Figure 4.**
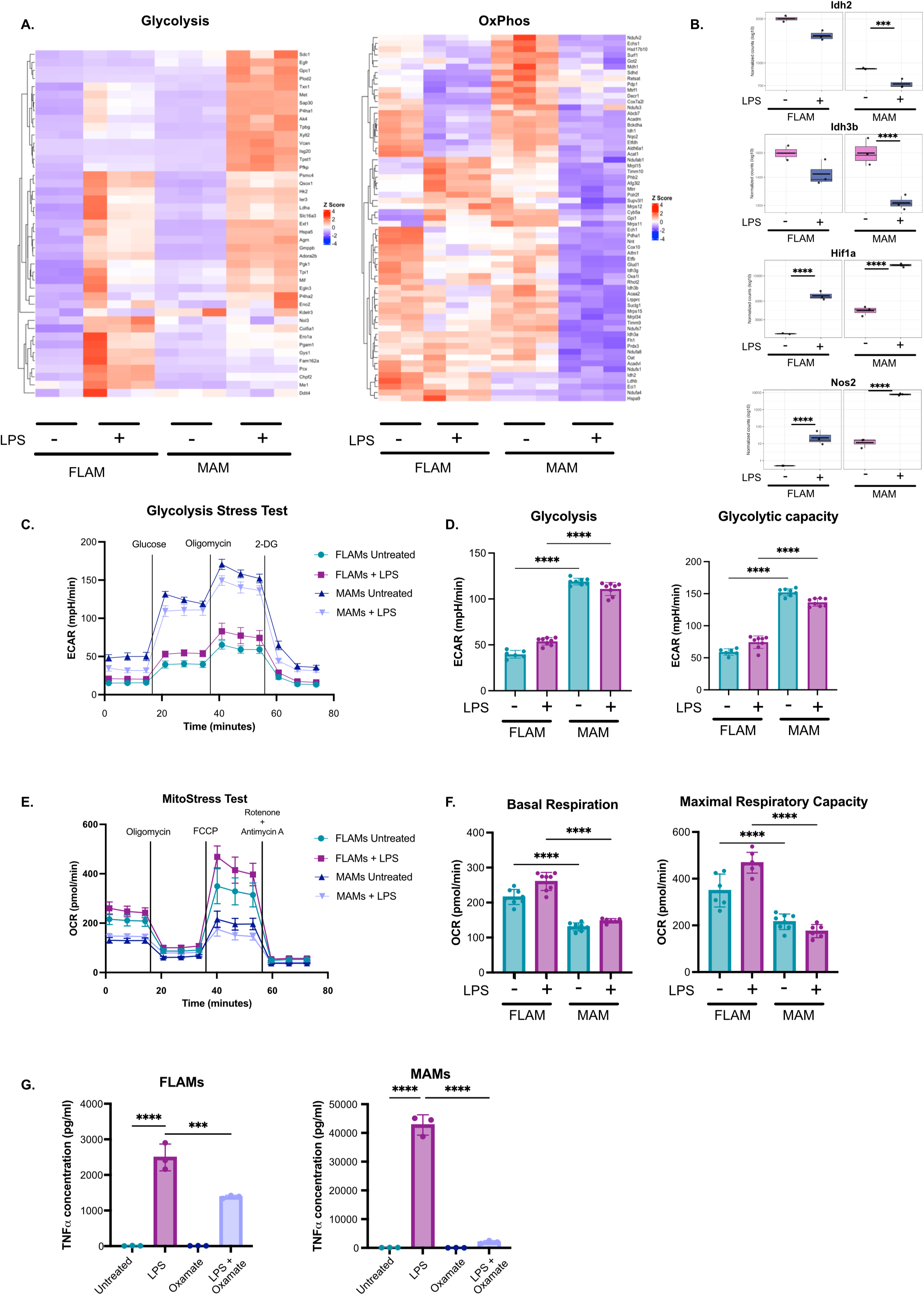
MAMs are highly dependent on glycolysis compared to FLAMs which contributes to increased inflammation. **A)** Normalized counts of glycolysis or oxidative phosphorylation genes from the KEGG pathway set were compared across samples of untreated or LPS treated MAMs and FLAMs. The color scale represents the z-score calculated from normalized read counts across samples for each gene. **B)** Normalized counts of a subset of glycolysis and OxPhos genes that were differentially regulated in FLAMs and MAMs with and without LPS treatment. ***p<.001 based on adjusted *p*-values using DESeq2 comparisons. **C)** The extracellular flux rate from a glycolysis stress test comparing untreated or LPS treated MAMs and FLAMs. **D)** The quantified glycolysis rate and glycolytic capacity were quantified from the experiment in F. Data in C and D are representative of three independent experiments ****p<.0001 by one-way ANOVA with a tukey test. **E)** The oxygen consumption rate was quantified during a mitostress test for untreated or LPS treated MAMs and FLAMs. **F)** The basal respiration and the maximum respiratory capacity were quantified from the Mitostress test in E. Data in E and F are representative of three independent experiments **** p<.0001 by one-way ANOVA with a tukey test. G) MAMs and FLAMs were pretreated with Oxamate followed by leaving cells untreated or activating with LPS. The next day supernatants were quantified for TNF by ELISA. Data are representative of three independent experiments ***p<.001 ****p<.0001 by one-way ANOVA with a tukey test.

To directly confirm the metabolic state of these cells, we next used a Seahorse analyzer to monitor the metabolic flux of both MAMs and FLAMs. First, we conducted a glycolysis stress on MAMs and FLAMs treated with and without LPS to determine the glycolytic capacity of these cells (Figure 4C). We observed that that MAMs have higher glycolysis rate, glycolytic capacity, compared to FLAMs under resting and LPS-activated conditions (Figure 4D). To define the respiratory capacity of these cells we next performed a Mito Stress test on MAMs and FLAMs that were untreated or treated with LPS (Figure 4E). We found that resting FLAMs have a significantly higher basal and maximal OCR compared to MAMs (Figure 4F). Surprisingly, this difference was further exacerbated by LPS treatment with FLAMs increasing their maximal OCR while MAMs maximal OCR decreased. Altogether these data support a model where MAMs drive high levels of glycolysis following LPS treatment while FLAMs stably maintain oxidative phosphorylation which correlates with their inflammatory profiles.

To understand the link between high glycolysis in MAMs and inflammatory cytokine production we next tested whether inhibition of glycolysis impacts the response to LPS. FLAMs and MAMs were left untreated or were pretreated with the lactate dehydrogenase inhibitor oxamate followed by stimulation with or without LPS. We then quantified the production of TNF by ELISA (Figure 4G). We observed that while oxamate treated FLAMs showed a modest but significant two-fold change in TNF following LPS stimulation, MAMs showed a massive 1000-fold decrease in TNF. These data show MAMs have high levels of glycolysis compared to FLAMs and these differences in metabolic regulation are critical to the inflammatory state of the cells.

### Driving lipid metabolism in MAMs does not reverse their hyperinflammatory response

We next were interested in understanding why FLAMs remained less inflammatory following LPS treatment and if we could activate pathways in MAMs to prevent their robust inflammatory responses. From our RNAseq dataset above, GSEA uncovered a robust enrichment of fatty acid metabolism genes in FLAMs compared to MAMs (Figure 5A). To directly examine how the expression of fatty acid metabolism is regulated in MAMs in FLAMs we compared the expression of genes associated with the Hallmark fatty acid metabolism pathway (Figure 5B). We found high expression of this pathway in FLAMs that was decreased following LPS. However, for MAMs, baseline expression of the fatty acid pathway matched LPS-treated FLAMs. Furthermore, we found that LPS treated MAMs further decreased the expression of these genes. When we plotted the normalized counts for a subset of these genes, we found that FLAMs maintain high expression of acetyl CoA acetyltransferase-2 (ACAT2), fatty acid synthase (FASN), and squalene epoxidase (SQLE) (Figure 5C). And these differences become more dramatic following LPS activation. To understand transcription factors that may play a role in this response we next examined a key regulator of lipid metabolism in AMs, the transcription factor Pparg. We noted that the expression of PPARg in FLAMs and MAMs was inversely correlated with the expression of the proinflammatory transcription factors NFKB1 and IRF7 (Figure 5D). Given the links between lipid metabolism and inflammation in MAMs and FLAMs we hypothesized that altering lipid metabolism would also modulate the inflammatory state of MAMs. To test this hypothesis, we compared the transcriptional responses of MAMs and FLAMs grown in the presence of two distinct lipid metabolism modulators, the PPARγ agonist rosiglitazone and natural calf lung surfactant, Infasurf®, which is metabolized by AMs in the lungs. MAMs or FLAMs were grown in the absence or presence of infasurf for one week or treated with rosiglitazone overnight prior to activation with or without LPS then bulk RNA sequencing was done (Supplementary Table 3). PCA analysis showed that FLAMs, regardless of condition, clustered closely together suggesting they are transcriptionally similar and not majorly impacted by either lipid treatment or LPS activation (Figure 5E). In contrast, there was a clear separation among the MAM samples that was only dependent on LPS activation. This suggests minimal effects of rosiglitazone and infasurf treatment on the overall MAM transcriptional profiles.

**Figure 5.**
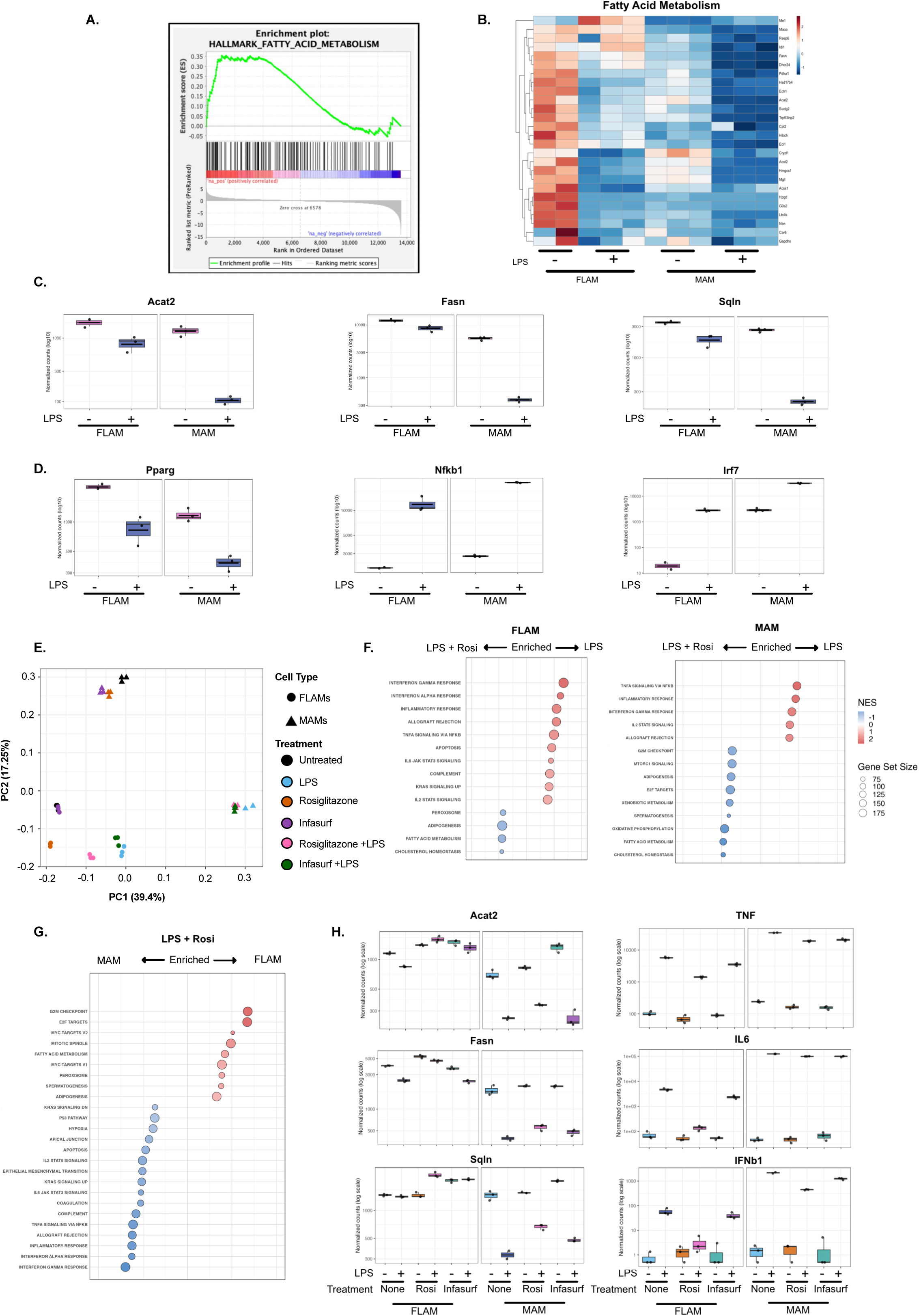
MAMs drive less robust fatty acid metabolism compared to FLAMs. **A)** Leading edge analysis of the Hallmark Fatty Acid Metabolism pathway from the RNAseq experiments presented in Figure 3 comparing MAMs and FLAMs in resting conditions. **B)** Normalized counts of the fatty acid metabolism Hallmark pathway were compared across samples of untreated or LPS treated MAMs and FLAMs. The color scale represents the z-score calculated from normalized read counts across samples for each gene. **C)** Normalized counts of a subset of fatty acid metabolism genes that were differentially regulated in FLAMs and MAMs with and without LPS treatment. ***p<.001 based on adjusted *p*-values using DESeq2 comparisons. **D)** Normalized counts of key transcription factors associated with lipid metabolism (Pparg) or inflammation (Nfkb1 and Irf7) that were differentially regulated in FLAMs and MAMs with and without LPS treatment. ***p<.001 based on adjusted *p*-values using DESeq2 comparisons. **E)** A principal component analysis (PCA) plot comparing the transcriptome of from MAMs and FLAMs that were untreated, grown in infasurf for 7 days, or treated with Rosiglitazone overnight following stimulation with or without LPS (12.5ng/ml) for 6 hours. **F)** Gene set enrichment analysis of pathways enriched in (Left) FLAMs or (Right) MAMs treated with LPS in the presence or absence of Rosiglitazone. **G)** Gene set enrichment analysis of pathways enriched in MAMs or FLAMs treated with LPS and Rosiglitazone. **H)** Normalized counts of key (Left) lipid metabolism genes or (Right) inflammatory cytokines from the RNA sequencing experiment presented in Figure 3E above. ***p<.001 based on adjusted *p*-values using DESeq2 comparisons.

To better understand how these treatments modulate broadly impact gene expression we examined the pathways that were enriched between LPS treated FLAMs with and without Rosiglitazone or MAMs with and without Rosiglitazone (Figure 5F). While Rosiglitazone treatment of both MAMs and FLAMs resulted in an enrichment of pathways related to fatty acid metabolism and cholesterol homeostasis, we noted an enrichment of peroxisome pathways only in treated FLAMs. Furthermore, Rosiglitazone treated FLAMs showed more dramatic differences in inflammatory pathways and Rosiglitazone treatment of MAMs did not alter the enrichment of the type I IFN pathway unlike what we observed in FLAMs. We further confirmed these findings when we directly compared the pathways enriched between MAMs and FLAMs that were LPS and Rosiglitazone treated (Figure 5G). We saw a continued enrichment of inflammatory pathways in MAMs and strong enrichment in fatty acid metabolism and peroxisomal pathways in FLAMs. Thus, it appears that Rosiglitazone treatment of FLAMs is unable to robustly induce fatty acid metabolism leading to a continued hyperinflammatory state that is reversed by Rosiglitazone treatment of FLAMs. When we directly examined the expression changes of Acat2, Fasn, and Sqln across all conditions we found that FLAMs maintain high expression of these genes which is further enhanced by rosiglitazone treatment while MAMs continued to repress the expression of these genes following LPS activation, regardless of condition (Figure 5H). Finally, we compared the expression of the proinflammatory cytokines TNF, IL6 and IFNβ. In FLAMs, while infasurf treatment had a limited impact on inflammatory cytokines following LPS activation, Rosiglitazone significantly repressed the expression of each cytokine. In contrast, neither infasurf nor Rosiglitazone treatment of LPS activated MAMs robustly impacted the expression of these cytokines which remained high in all LPS conditions. Together, these data show that while FLAMs can be modulated by the activity of rosiglitazone, MAMs are more recalcitrant to its effects on inflammation.

### MAMs remain hyperinflammatory within the lung environment in a model of acute lung injury

Fully defining differences between MAMs and FLAMs requires an *in vivo* approach to understand their role in the lungs. To address this critical question, we next optimized a FLAM/MAM transfer model into mice deficient in the GM-CSFR. Due to the lack of functional GM-CSF signaling, these mice do not have tissue resident AMs and develop severe pulmonary alveolar proteinosis (**PAPs**) which is driven by the buildup of non-metabolized surfactant. As a first step we transferred either FLAMs, MAMs, or PBS intranasally into GM-CSFR-/- mice and 1 week later we isolated bronchial lavage fluid (**BAL**). We reasoned that since FLAMs and MAMs have a functional GM-CSFR they would be capable of being maintained in the lungs unlike any endogenous cells from the animal. In GM-CSFR-/- mice the buildup of surfactant is known to increase the turbidity of the BAL and thus we measured the optical density of the BAL across each condition. While BAL from mice that received PBS had a high OD, we noted a greater than five-fold decrease in mice that received either FLAMs or MAMs (Figure 6A). When we analyzed the BAL by flow cytometry, we noted a robust alveolar macrophage population in mice that received either FLAMs or MAMs but not PBS (Figure 6B and 6C). Thus, both MAMs and FLAMs metabolize surfactant and can be maintained long-term in the lung environment following transfer into GM-CSFR mice that lack AMs. Due to their hyperinflammatory state we next hypothesized that mice receiving MAMs would drive more inflammation in the lungs following administration of LPS. To test this hypothesis we transferred MAMs, FLAMs, or PBS into GM-CSFR KO mice and 4 weeks later we instilled 1mg/kg LPS intratracheally. 24 hours later we isolated BAL and quantified the production of TNF, IL6 and the IFN stimulated gene CXCL10. We observed a significant increase in each cytokine in mice that received MAMs compared to FLAMs (Figure 6D). Thus, MAMs not only recapitulate phenotypes associated with myeloid-derived AMs *ex vivo*, but also within the lung environment.

**Figure 6.**
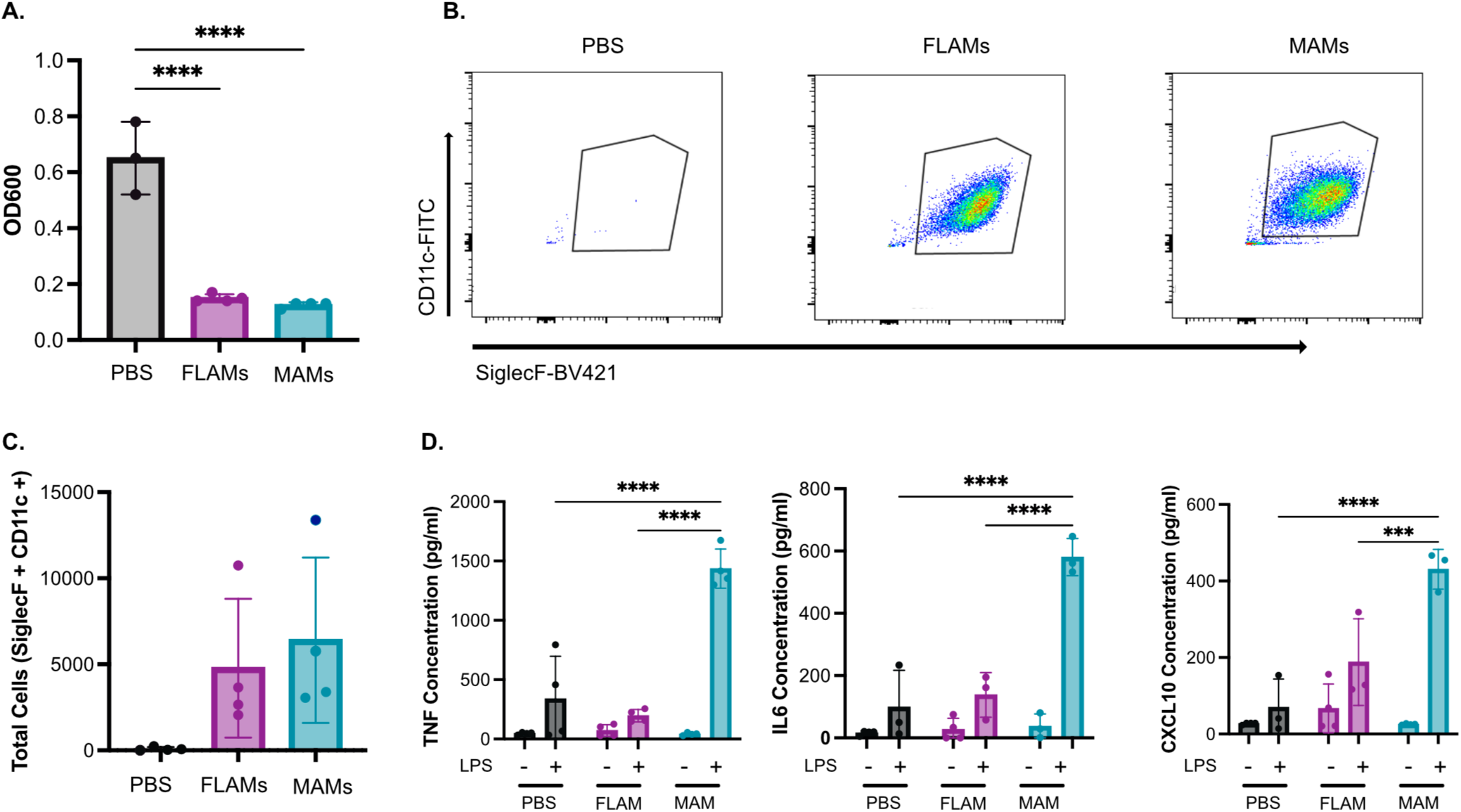
MAMs function *in vivo* and retain their hyperinflammatory response. BAL was isolated from GM-CSFR mice following transfer of PBS, FLAMs, and MAMs for 4 weeks **A)** The optical density at 600nm was determined from each sample. ****p<.0001 by one-way ANOVA with a Tukey test. **B)** Representative flow cytometry plot examining SiglecF+ CD11c+ cells. **C)** Quantification of total SiglecF+ CD11c+ cells. **p<.01 by one-way ANOVA with a tukey test. Data in A-C are representative of 3 independent experiments. **D)** PBS, FLAMs, and MAMs were transferred into GM-CSFR mice for 4 weeks then challenged with PBS or LPS instilled by oropharyngeal route. The following day BAL was isolated, and the indicated cytokines were quantified by ELISA. Data are representative of two independent experiments. ***p<.001 ****p<.0001 by one-way ANOVA with a tukey test.

## Discussion

Regulation of pulmonary inflammation is critically important to maintain lung function while also properly removing potential threats inhaled from the air^33–35^. While macrophages are key mediators of the initial inflammatory state in the lungs, how distinct macrophage subsets of different ontogeny regulate their inflammation and contribute to pulmonary disease remains unclear^10,36,37^. To address this key gap in knowledge, here we developed a model of myeloid-derived alveolar macrophages, MAMs, that enables us to compare and contrast the underlying inflammatory control to a tissue resident AMs or a model of AMs we developed previously, FLAMs ^13,25,28^. Importantly, our results show that MAMs recapitulate many phenotypes associated with myeloid-derived AMs both *ex vivo* and *in vivo ^12,24,38^*. Leveraging this powerful approach, we began dissecting the underlying mechanisms that drive key inflammatory regulation pathways in both MAMs and FLAMs. We found that many differences between MAMs and FLAMs including inflammation, epigenetic regulation, and metabolism are all hardwired based on ontogeny. Altogether our data show that MAMs will be an important model to understand the regulation of distinct pulmonary macrophage subsets and dissect their contribution of pulmonary disease *in vivo*.

Myeloid-derived AMs are increasingly acknowledged as a key mediator of important lung pathologies including post influenza bacterial pneumonia^10,39^. In this condition, a primary influenza infection depletes tissue resident AMs shifting the balance in the lungs towards myeloid derived-AMs^39,40^. Following the sensing of a secondary bacterial infection by these myeloid-derived AMs, there is increased inflammatory responses, immune cells recruitment and tissue damage resulting in severe disease^10,40^. However, our mechanistic understanding of myeloid-derived AMs is limited due to the requirement for *in vivo* studies. MAMs overcomes this limitation and key consideration of their utility is the similarity of MAMs to myeloid-derived AMs in the lungs. Our data show that MAMs recapitulate key phenotypes associated with Myeloid-derived AMs *in vivo*. MAMs are more inflammatory than fetal-liver derived AMs producing higher levels of distinct cytokines including TNF, IL1, IL6 and IFNβ. MAMs are also epigenetically distinct showing key differences in DNA methylation from fetal liver derived AMs. These differences align with those described in the lungs between tissue resident AMs and myeloid-derived AMs^10,12^. Importantly, at baseline conditions we found that MAMs and FLAMs both transcriptionally and functionally are similar. Only following activation with LPS are the underlying differences between MAMs and FLAMs robustly observed, similar to what is seen *in vivo*. Future work will be needed to understand how interactions with pathogens differs between MAMs and FLAMs. A range of bacterial pathogens including *Klebsiella pneumoniae*, *Streptococcus pneumonia*, and *Staphylococcus aureus* all are implicated in post-influenza bacterial pneumoniae, but a detailed understanding of host-pathogen interactions with distinct alveolar macrophage subsets remains to be completed^10,41,42^. These important gaps in knowledge can now begin to be filled by using MAMs as to understand the genetic regulation of this critically important lung macrophage subset.

One of the most striking differences between MAMs and FLAMs is their metabolic regulation. FLAMs are highly dependent on lipid metabolism and OxPhos and do not robustly shift their metabolism following activation with LPS ^25,28^. In contrast, MAMs were highly glycolytic both in resting and activated conditions. This glycolytic dependency drove their inflammatory state as blocking glycolysis reduced their production of cytokines. What drives these underlying differences in metabolism remains to be understood. One possible contributor that drives metabolic differences between MAMs and FLAMs are peroxisomes and their dependency on lipid metabolism. Peroxisomes are highly expressed in both FLAMs and tissue resident AMs, and the loss of peroxisomes in these cells has dramatic impacts on their function and survival in the lungs ^27,29^. Our data also suggest that modulating peroxisomes through the activity of rosiglitazone or the addition of lipid surfactants, further modulates the inflammatory state of FLAMs^43^. In contrast, rosiglitazone and infasurf had limited effects on MAMs and their inflammatory state. While this suggests underlying differences in lipid metabolism drive differences between MAMs and FLAMs, several open questions remain. It remains to be understood what drives distinct metabolic states in these cells and a careful understanding of peroxisome dynamics with MAMs and FLAMs at baseline and following activation. Future work using peroxisome deficient mice or other peroxisome activators such as 4-PBA will help to better understand the important role these organelles play in the underlying functions of distinct alveolar macrophages subsets.

Our finding that both MAMs and FLAMs function in the lung environment following transplantation is an important step to fully characterize the role of these cells *in vivo*. By leveraging the AM-deficiency and surfactant buildup in GM-CSFR-/- mice, we found that both MAMs and FLAMs were equally capable of metabolizing surfactant and being maintained long term. The ability to control the population of alveolar macrophages in the lungs with high precision can now be leveraged to understand how the balance between distinct alveolar macrophage subsets contributes to tissue homeostasis and responses to infection. Not only can we transfer either FLAMs or MAMs into the lungs, but we can also mix these populations to define their interactions and responses in a more complex lung environment. The only current strategy to study myeloid-derived AMs is by infecting mice with a pathogen, which is likely to change more within the lung environment than just the macrophage populations. The MAM transfer model provides a unique approach to understand how the arrival of myeloid-derived AMs into the lungs directly modulates downstream host responses and tissue damage. More importantly, a reproducible *ex vivo* MAM system will enable the development of functional genetic tools similar to what we optimized previously FLAMs. Combining genetic approaches in MAMs with *in vivo* transfers will position us to dissect pathways that are required for the myeloid-derived AM inflammatory state and function within the lung environment.

An important consideration with pulmonary disease states is identifying potential pathways that can be targeted to modulate inflammatory and pathologic outcomes. Our data which is in line with published studies, clearly show that metabolic networks are key to these inflammatory states. However, simply activating lipid metabolic pathways in MAMs is insufficient to modulate their inflammation. Thus, future studies will be positioned to understand pathways that are required for myeloid-derived AM inflammation that can be targeted therapeutically. This important advance will allow us to make inroads to help patients susceptible to pulmonary disease states associated with inflammation and disrupted balance in the lung macrophage populations.

## Acknowledgements

This work was supported by grants from the National Institutes of Health: and R35 GM146795 (A.J.O.).The Attune CytPix, located in the MSU Flow Cytometry Core Facility, is supported by the Equipment Grants Program, award #2022-70410-38419, from the U.S. Department of Agriculture (USDA). The authors wish to thank the Olive Lab for helpful discussions and feedback on data and the manuscript. The author would like to thank the Michigan State University Center for Advanced Microscopy Core Facility (RRID:SCR_027702) for assistance with scanning electron microscopy.

## Figure Legends

**Supplementary Table 1.** CpG beta-values comparing MAMs and FLAMs with and without LPS

**Supplementary Table 2.** Normalized count table for MAMs and FLAMs +/- LPS treatment

**Supplementary Table 3.** Normalized count table for MAMs and FLAMs with Rosiglitazone, Infasurf and LPS.

## References

1. McQuattie-Pimentel AC, Ren Z, Joshi N, et al. The lung microenvironment shapes a dysfunctional response of alveolar macrophages in aging. J Clin Invest. 2021;131(4). doi:10.1172/JCI140299

2. Byrne AJ, Mathie SA, Gregory LG, Lloyd CM. Pulmonary macrophages: key players in the innate defence of the airways. Thorax. 2015;70(12):1189. doi:10.1136/thoraxjnl-2015-207020

3. Mass E, Ballesteros I, Farlik M, et al. Specification of tissue-resident macrophages during organogenesis. Science (1979). 2016;353(6304):aaf4238. doi:10.1126/science.aaf4238

4. Davies LC, Jenkins SJ, Allen JE, Taylor PR. Tissue-resident macrophages. Nat Immunol. 2013;14(10):986-995. doi:10.1038/ni.2705

5. Guilliams M, De Kleer I, Henri S, et al. Alveolar macrophages develop from fetal monocytes that differentiate into long-lived cells in the first week of life via GM-CSF. Journal of Experimental Medicine. 2013;210(10):1977–1992. doi:10.1084/jem.20131199

6. Rosseau S, Hammerl P, Maus U, et al. Phenotypic Characterization of Alveolar Monocyte Recruitment in Acute Respiratory Distress Syndrome. 2000. http://www.ajplung.org

7. Shibata Y, Berclaz PY, Chroneos ZC, Yoshida M, Whitsett JA, Trapnell BC. GM-CSF Regulates Alveolar Macrophage Differentiation and Innate Immunity in the Lung through PU.1. Immunity. 2001;15(4):557–567. 10.1016/S1074-7613(01)00218-7

8. Yu X, Buttgereit A, Lelios I, et al. The Cytokine TGF-β Promotes the Development and Homeostasis of Alveolar Macrophages. Immunity. 2017;47(5):903–912.e4. doi:10.1016/j.immuni.2017.10.007

9. Yu X, Buttgereit A, Lelios I, et al. The Cytokine TGF-β Promotes the Development and Homeostasis of Alveolar Macrophages. Immunity. 2017;47(5):903–912.e4. doi:10.1016/j.immuni.2017.10.007

10. Aegerter H, Kulikauskaite J, Crotta S, et al. Influenza-induced monocyte-derived alveolar macrophages confer prolonged antibacterial protection. Nat Immunol. 2020;21(2):145–157. doi:10.1038/s41590-019-0568-x

11. Pahari S, Arnett E, Simper J, et al. A new tractable method for generating human alveolar macrophage-like cells in vitro to study lung inflammatory processes and diseases. mBio. 2023;14(4):e0083423. doi:10.1128/mbio.00834-23

12. Gibbings SL, Goyal R, Desch AN, et al. Transcriptome analysis highlights the conserved difference between embryonic and postnatal-derived alveolar macrophages. Blood. 2015;126(11):1357–1366. doi:10.1182/blood-2015-01-624809

13. Gilliland HN, Soverina S, Conner KN, Vielma TE, Olive AJ. Differential activation of NF-κB and HIF-1α between alveolar-like macrophages and myeloid-derived macrophages drive inflammatory differences following *Mycobacterium abscessus* infection. bioRxiv. Published online January 1, 2025:2025.04.03.647026. doi:10.1101/2025.04.03.647026

14. Jia L, Luo H, Li L, et al. Targeting complement hyperactivation: a novel therapeutic approach for severe pneumonia induced by influenza virus/staphylococcus aureus coinfection. Signal Transduct Target Ther. 2023;8(1):467. doi:10.1038/s41392-023-01714-y

15. Kawai T, Ikegawa M, Ori D, Akira S. Decoding Toll-like receptors: Recent insights and perspectives in innate immunity. Immunity. 2024;57(4):649–673. doi:10.1016/j.immuni.2024.03.004

16. Chen R, Zou J, Chen J, Zhong X, Kang R, Tang D. Pattern recognition receptors: function, regulation and therapeutic potential. Signal Transduct Target Ther. 2025;10(1):216. doi:10.1038/s41392-025-02264-1

17. Watts C. Location, location, location: identifying the neighborhoods of LPS signaling. Nat Immunol. 2008;9(4):343–345. doi:10.1038/ni0408-343

18. Ciesielska A, Matyjek M, Kwiatkowska K. TLR4 and CD14 trafficking and its influence on LPS-induced pro-inflammatory signaling. Cellular and Molecular Life Sciences. Springer Science and Business Media Deutschland GmbH. 2021;78(4):1233–1261. doi:10.1007/s00018-020-03656-y

19. Zhang J, Chang J, Beg MA, et al. Na/K-ATPase suppresses LPS-induced pro-inflammatory signaling through Lyn. iScience. 2022;25(9). doi:10.1016/j.isci.2022.104963

20. Olson GS, Murray TA, Jahn AN, et al. Type I interferon decreases macrophage energy metabolism during mycobacterial infection. Cell Rep. 2021;35(9). doi:10.1016/j.celrep.2021.109195

21. Viola A, Munari F, Sánchez-Rodríguez R, Scolaro T, Castegna A. The metabolic signature of macrophage responses. Front Immunol. Frontiers Media S.A. 2019;10(JULY). doi:10.3389/fimmu.2019.01462

22. O’Neill LAJ, Kishton RJ, Rathmell J. A guide to immunometabolism for immunologists. Nat Rev Immunol. Nature Publishing Group. 2016;16(9):553–565. doi:10.1038/nri.2016.70

23. Hackett EE, Charles-Messance H, O’Leary SM, et al. Mycobacterium tuberculosis Limits Host Glycolysis and IL-1β by Restriction of PFK-M via MicroRNA-21. Cell Rep. 2020;30(1):124–136.e4. doi:10.1016/j.celrep.2019.12.015

24. Gorki AD, Symmank D, Zahalka S, et al. Murine Ex Vivo Cultured Alveolar Macrophages Provide a Novel Tool to Study Tissue-Resident Macrophage Behavior and Function. Am J Respir Cell Mol Biol. 2022;66(1):64–75. doi:10.1165/rcmb.2021-0190OC

25. Thomas ST, Wierenga KA, Pestka JJ, Olive AJ. Fetal Liver–Derived Alveolar-like Macrophages: A Self-Replicating Ex Vivo Model of Alveolar Macrophages for Functional Genetic Studies. Immunohorizons. 2022;6(2):156–169. doi:10.4049/immunohorizons.2200011

26. Schneider C, Nobs SP, Kurrer M, Rehrauer H, Thiele C, Kopf M. Induction of the nuclear receptor PPAR-γ 3 by the cytokine GM-CSF is critical for the differentiation of fetal monocytes into alveolar macrophages. Nat Immunol. 2014;15(11):1026–1037. doi:10.1038/ni.3005

27. Wei X, Qian W, Narasimhan H, et al. Macrophage peroxisomes guide alveolar regeneration and limit SARS-CoV-2 tissue sequelae. Science (1979). 2025;387(6738). doi:10.1126/science.adq2509

28. Thomas SM, McGee AP, Vielma TE, et al. TGFβ primes alveolar-like macrophages to induce type I IFN following TLR2 activation. The Journal of Immunology. 2026;215(5):vkag090. doi:10.1093/jimmun/vkag090

29. Kim J, Bai H. Peroxisomal Stress Response and Inter-Organelle Communication in Cellular Homeostasis and Aging. Antioxidants. MDPI. 2022;11(2). doi:10.3390/antiox11020192

30. Gorki AD, Symmank D, Zahalka S, et al. Murine Ex Vivo Cultured Alveolar Macrophages Provide a Novel Tool to Study Tissue-Resident Macrophage Behavior and Function. Am J Respir Cell Mol Biol. 2022;66(1):64–75. doi:10.1165/rcmb.2021-0190OC

31. Thomas ST, Wierenga KA, Pestka JJ, Olive AJ. Fetal Liver–Derived Alveolar-like Macrophages: A Self-Replicating Ex Vivo Model of Alveolar Macrophages for Functional Genetic Studies. Immunohorizons. 2022;6(2):156–169. doi:10.4049/immunohorizons.2200011

32. Nielsen TB, Yan J, Luna B, Spellberg B. Murine Oropharyngeal Aspiration Model of Ventilator-associated and Hospital-acquired Bacterial Pneumonia. JoVE. 2018;(136):e57672. doi:doi:10.3791/57672

33. Branchett WJ, Lloyd CM. Regulatory cytokine function in the respiratory tract. Mucosal Immunol. 2019;12(3):589–600. doi:10.1038/s41385-019-0158-0

34. Kopf M, Schneider C, Nobs SP. The development and function of lung-resident macrophages and dendritic cells. Nat Immunol. 2015;16(1):36–44. doi:10.1038/ni.3052

35. Hsia CCW, Hyde DM, Weibel ER. Lung Structure and the Intrinsic Challenges of Gas Exchange. Compr Physiol. 2016;6(2):827–895. 10.1002/j.2040-4603.2016.tb00698.x

36. Huang L, Nazarova E V, Tan S, Liu Y, Russell DG. Growth of Mycobacterium tuberculosis in vivo segregates with host macrophage metabolism and ontogeny. Journal of Experimental Medicine. 2018;215(4):1135–1152. doi:10.1084/jem.20172020

37. Pisu D, Huang L, Grenier JK, Russell DG. Dual RNA-Seq of Mtb-Infected Macrophages In Vivo Reveals Ontologically Distinct Host-Pathogen Interactions. Cell Rep. 2020;30(2):335–350.e4. 10.1016/j.celrep.2019.12.033

38. Bragazzi Cunha J, Leix K, Sherman EJ, et al. Type I interferon signaling induces a delayed antiproliferative response in respiratory epithelial cells during SARS-CoV-2 infection. J Virol. 2023;97(12). doi:10.1128/jvi.01276-23

39. Hou F, Xiao K, Tang L, Xie L. Diversity of Macrophages in Lung Homeostasis and Diseases. Front Immunol. Frontiers Media S.A. 2021;12. doi:10.3389/fimmu.2021.753940

40. Morris DE, Cleary DW, Clarke SC. Secondary bacterial infections associated with influenza pandemics. Front Microbiol. Frontiers Media S.A. 2017;8(JUN). doi:10.3389/fmicb.2017.01041

41. Liu T, Miller LM, Dresden BP, et al. Influenza virus and Staphylococcus aureus super-infection disrupts spatially coordinated cellular immunity in the mouse lung. Commun Biol. 2025;8(1):1837. doi:10.1038/s42003-025-09268-1

42. Soverina S, Gilliland HN, Olive AJ. Pathogenicity and virulence of Mycobacterium abscessus. Virulence. 2025;16(1):2508813. doi:10.1080/21505594.2025.2508813

43. Christofides A, Konstantinidou E, Jani C, Boussiotis VA. The role of peroxisome proliferator-activated receptors (PPAR) in immune responses. Metabolism. W.B. Saunders. 2021;114. doi:10.1016/j.metabol.2020.154338

